# Educational Attainment Polygenic Scores, Socioeconomic Factors, and Cortical Structure in Children and Adolescents

**DOI:** 10.1101/2021.09.17.460810

**Authors:** Emily C. Merz, Jordan Strack, Hailee Hurtado, Uku Vainik, Michael Thomas, Alan Evans, Budhachandra Khundrakpam

**Affiliations:** Department of Psychology, Colorado State University; University of Tartu, Estonia; Montreal Neurological Institute, McGill University

## Abstract

**Background:** Genome-wide polygenic scores for educational attainment (PGS-EA) and socioeconomic factors, which are correlated with each other, have been consistently associated with academic achievement and general cognitive ability in children and adolescents. Yet, the independent associations of PGS-EA and socioeconomic factors with specific underlying factors at the neural and neurocognitive levels are not well understood. The goal of this study was to examine the unique contributions of PGS-EA and parental education to cortical thickness (CT), cortical surface area (SA), and neurocognitive skills in children and adolescents.

**Methods:** Participants were typically developing children and adolescents (3-21 years of age; 53% male; *N* = 391). High-resolution, T1-weighted magnetic resonance imaging data were acquired. PGS-EA were computed based on the most recent genome-wide association study of educational attainment. Sustained attention, inhibitory control, working memory, vocabulary, and episodic memory were measured.

**Results:** PGS-EA and parental education were independently and significantly associated with SA, vocabulary, and attention outcomes but were not associated with CT. Vertex-wise analyses indicated that higher PGS-EA was significantly associated with greater SA in the left medial orbitofrontal gyrus and inferior frontal gyrus after accounting for parental education. Higher parental education was significantly associated with greater SA in the left parahippocampal gyrus after accounting for PGS-EA.

**Conclusions:** These findings suggest that education-linked genetics may influence SA, particularly in certain frontal regions, leading to variability in academic achievement. Results suggested genetic confounding in associations between parental education and SA in children and adolescents, with these associations remaining significant after controlling for PGS-EA.

---

Elucidating how genetic and environmental factors influence brain development in children and adolescents is an important task for researchers. Recent scientific advances allowing the computation of genome-wide polygenic scores have led to ground-breaking insights into genetic effects on cognitive and health outcomes (Armstrong-Carter et al., 2021; Plomin & von Stumm, 2018). Polygenic scores are derived using genome-wide association studies (GWAS) by aggregating the contributions of all known genetic variants associated with the phenotype of interest (Plomin & von Stumm, 2018). GWAS have identified genetic variants robustly associated with educational attainment (years of education) and yielded genome-wide polygenic scores for educational attainment (PGS-EA) that significantly predict years of education (Lee et al., 2018; Okbay et al., 2016; Rietveld et al., 2013), academic achievement (Selzam et al., 2017; von Stumm et al., 2020; Ward et al., 2014), and general cognitive ability (Allegrini et al., 2019; Belsky et al., 2016; Judd et al., 2020; Wertz et al., 2018) in independent samples. However, the associations between PGS-EA and the underlying factors at the neural and neurocognitive levels in children and adolescents are not well understood.

Building from decades of research demonstrating socioeconomic disparities in cognitive development (McLoyd, 1998), recent studies have shed light on the neural mechanisms underlying these associations (Farah, 2017). Socioeconomic factors, such as parental education and family income, have been repeatedly associated with brain structure in children and adolescents (Farah, 2017; McDermott et al., 2019; Noble et al., 2015; Noble & Giebler, 2020), with evidence pointing to the environmental factors involved in these associations (Merz et al., 2020). Yet, the environments in which children are raised are associated with the genotypes they inherit from their parents (i.e., gene-environment correlation) (Plomin et al., 2016). In one example of a passive gene-environment correlation, more educated parents provide both a genetic propensity for higher educational attainment and cognitively stimulating home environments to their children. Indeed, the associations between socioeconomic factors and children’s academic achievement may be partially attributable to genetic transmission (Belsky et al., 2016, 2018; Krapohl & Plomin, 2016; von Stumm et al., 2020). However, the unique role of socioeconomic factors in predicting the underlying neural and neurocognitive measures independent of genetic factors is not well understood. As such, the goal of this study was to examine the independent associations of PGS-EA and parental education with cortical structure and neurocognitive skills in children and adolescents.

## PGS-EA and Cortical Structure

In recent years, researchers have leveraged GWAS techniques to investigate the genetics of educational attainment (Lee et al., 2018; Okbay et al., 2016; Rietveld et al., 2013). Educational attainment is a demographic measure collected in most studies, allowing large studies to be conducted on this phenotype. The most recent GWAS (EA3) included data from over a million adults of European ancestry and identified 1,271 significant single-nucleotide polymorphisms (SNPs) (Lee et al., 2018). A polygenic score derived from the results explained up to 13% of the variance in educational attainment in independent samples (Lee et al., 2018).

To our knowledge, only two neuroimaging studies to date have focused on PGS-EA and cortical structure in children and adolescents. In one study, PGS-EA were significantly positively associated with total brain volume in a large sample of 10-year-olds (Alemany et al., 2019). Cortical volume is a composite of cortical surface area (SA) and cortical thickness (CT), which are genetically, developmentally, and phenotypically independent (Panizzon et al., 2009; Raznahan et al., 2011; Winkler et al., 2010). In a study that examined SA and CT separately, PGS-EA were significantly positively associated with global SA but not significantly associated with global CT in adolescents (Judd et al., 2020). In addition, PGS-EA were significantly associated with regional SA in the right intraparietal sulcus (Judd et al., 2020).

## Socioeconomic Factors and Cortical Structure

Numerous studies have revealed associations between socioeconomic factors and cortical structure in children and adolescents (Farah, 2017; Noble & Giebler, 2020). In these studies, higher parental education and family income have been significantly associated with greater SA (Judd et al., 2020; McDermott et al., 2019; Noble et al., 2015) and CT (Lawson et al., 2013; Mackey et al., 2015; McDermott et al., 2019; Romeo et al., 2017). These socioeconomic differences in cortical structure have been found to be most prominent in frontal and temporal regions crucial to language, executive function, and memory (McDermott et al., 2019; Noble et al., 2015). Yet, to our knowledge, only one study has examined associations between socioeconomic factors and cortical structure while controlling for PGS-EA in children and adolescents. In this study, parental education remained significantly positively associated with total SA after controlling for PGS-EA in adolescents (Judd et al., 2020).

## PGS-EA, Socioeconomic Factors, and Neurocognitive Measures

Although multiple studies have demonstrated associations between PGS-EA and general cognitive ability (Plomin & von Stumm, 2018), a smaller body of work has shown associations between PGS-EA and specific neurocognitive skills that underlie general cognitive ability. PGS-EA has been significantly associated with vocabulary, executive function (inhibitory control, working memory), and episodic memory in children and adolescents (Domingue et al., 2015; Judd et al., 2020; Loughnan et al., 2021; Rea-Sandin et al., 2021). In a largely separate literature, greater family income and parental education have been significantly associated with higher levels of these neurocognitive skills (Lawson et al., 2017; Merz et al., 2019; Noble et al., 2005, 2007). However, the independent associations of PGS-EA and socioeconomic factors with neurocognitive skills in children and adolescents are not well understood.

## Current Study

The goal of this study was to examine the independent associations of PGS-EA and parental education with cortical structure and specific neurocognitive skills in children and adolescents. Participants were typically developing children and adolescents who ranged in age from 3-21 years (*N* = 391). High-resolution, T1-weighted MRI data were acquired, and attention, vocabulary, inhibitory control, working memory, and episodic memory were measured. PGS-EA were computed using results from the most recent GWAS of educational attainment (Lee et al., 2018). We conducted analyses of global measures of CT and SA and vertex-wise analyses of regional CT and SA. Parental education and family income were examined separately (rather than combined into an SES composite) because they have been associated with distinct aspects of children’s environments and relate differentially to children’s development (Duncan & Magnuson, 2012). While the main analyses focus on parental education, results for family income are presented in the supplemental material.

We hypothesized that PGS-EA and parental education would be independently associated with SA and possibly CT. Based on previous research (McDermott et al., 2019; Mitchell et al., 2020; Noble et al., 2015), we expected these associations to be most pronounced in frontal and temporal cortical regions. We also hypothesized that both PGS-EA and parental education would be associated with a range of neurocognitive skills.

Some research has suggested gene-by-SES interactions may predict cognitive ability such that the heritability of intelligence is lower in low-SES family environments and higher in high-SES family environments (Tucker-Drob & Bates, 2016). Thus, we also examined interactions between socioeconomic factors and PGS-EA in predicting cortical structure.

## Methods

### Participants

Data were obtained from the Pediatric Imaging, Neurocognition and Genetics (PING) study (http://ping.chd.ucsd.edu/) (Jernigan et al., 2016). The PING study is a large-scale, publicly available data set for investigating neuroimaging, cognition and genetics in typically-developing children and adolescents (Jernigan et al., 2016). Exclusionary criteria in the PING study included neurological disorders; history of head trauma; preterm birth; diagnosis of an autism spectrum disorder, bipolar disorder, schizophrenia, or significant intellectual disability; and contraindications for MRI (Jernigan et al., 2016).

In total, the PING study included cross-sectional data collected from nine different sites across the United States. Participants in the current study ranged from 3-21 years of age (*M* = 11.53, *SD* = 4.82) and were 53% male. Family income ranged from $4,500-$325,000 (*M* = 121,290.35, *SD* = 76,743.49); parental education ranged from 8-18 years (*M* = 15.73; *SD* = 1.86).

Written informed consent was provided by parents for all participants younger than 18 years of age and by the participants themselves if they were 18 years or older. Child assent was obtained for 7- to 17-year-old participants. Each site’s Institutional Review Board approved the study.

### Socioeconomic Factors

Educational attainment was averaged across parents. Both education and income data were originally collected in bins, which were recoded as the means of the bins for analysis, following from previous work (Noble et al., 2015). Family income was log-transformed to correct for positive skew. Family income and parental education were significantly correlated, *r* = .56, *p* < .0001.

### Genomic Data

The PING dataset includes 550,000 SNPs genotyped from saliva samples using Illumina Human660W-Quad BeadChip. Computation of polygenic scores followed steps similar to that of our previous study (Khundrakpam, Vainik, et al., 2020). Steps included preparation of the data for imputation using the “imputePrepSanger” pipeline (https://hub.docker.com/r/eauforest/imputeprepsanger/) and implemented on CBRAIN (Sherif et al., 2014) using Human660W-Quad_v1_A-b37-strand chip as reference. The next step involved data imputation with Sanger Imputation Service (McCarthy et al., 2016) using default settings and the Haplotype Reference Consortium, HRC (http://www.haplotype-reference-consortium.org/) as the reference panel. Using Plink 1.9 (Chang et al., 2015), the imputed SNPs were then filtered with the inclusion criteria: SNPs with unique names, only ACTG, and MAF > 0.05. All SNPs that were included had INFO scores R2 > 0.9 with Plink 2.0. Next, using polygenic score software PRSice 2.1.2 (Euesden et al., 2015) additional ambiguous variants were excluded, resulting in 4,696,385 variants being available for polygenic scoring. We filtered individuals with 0.95 loadings to the European principal component (GAF_Europe variable provided with the PING data), resulting in 526 participants. These participants were then used to compute 10 principal components with Plink 1.9. Polygenic scores based on the EA3 GWAS (Lee et al., 2018) were used in analyses. We clumped the data as per PRSice default settings (clumping distance = 250 kb, threshold r^2^ = 0.1).

PGS-EA were computed at different *p*-value thresholds and the most predictive one was chosen, following previous studies (Alemany et al., 2019; Du Rietz et al., 2018; Judd et al., 2020). The *p*-value threshold of *p* < 1 x 10^-7^ best explained variance in SA. Thus, this conservative *p*-value threshold was used for the main analyses. After matching with available variants in the data, this PGS-EA was based on 694 variants (see Table S1). Results for other *p*-value thresholds were consistent with the results reported.

### Image Acquisition and Preprocessing

Each site administered a standardized structural MRI protocol. Imaging data were collected using 3-Tesla scanners manufactured by General Electric, Siemens, and Philips. The imaging protocols and pulse sequence parameters used in the PING study have been published previously (Jernigan et al., 2016; Merz et al., 2018; Noble et al., 2015). T1-weighted images were acquired using a standardized high-resolution 3D RF-spoiled gradient echo sequence (Jernigan et al., 2016).

The raw T1-weighted imaging data for the PING study are publicly shared (https://nda.nih.gov/) for a subset of the sample (*n* = 934). The only difference between the full PING sample and the subsample with raw T1-weighted imaging data was that the full PING sample was older on average than the subsample (Khundrakpam, Choudhury, et al., 2020). We used the CIVET processing pipeline (https://mcin.ca/technology/civet/) developed at the Montreal Neurological Institute to compute CT measurements at 81,924 regions covering the entire cortex. Processing steps included non-uniformity correction of the T1-weighted image and then linear registration to the Talairach-like MNI152 template (created with the ICBM152 dataset). After repeating the non-uniformity correction using the template mask, the non-linear registration from the resultant volume to the MNI152 template is computed, and the transform used to provide priors to segment the image into gray matter, white matter, and cerebrospinal fluid. Inner and outer gray matter surfaces are then extracted using the Constrained Laplacian-based Automated Segmentation with Proximities (CLASP) algorithm. CT is then measured in native space using the linked distance between the two surfaces at 81,924 vertices. To impose a normal distribution on the corticometric data and increase the signal to noise ratio, each individual’s CT map was blurred using a 30-millimeter full width at half maximum surface-based diffusion smoothing kernel. Two independent reviewers performed quality control (QC) of the data, and only scans with consensus of the two reviewers were used. Exclusion criteria for QC procedure included: data with motion artifacts, a low signal to noise ratio, artifacts due to hyperintensities from blood vessels, surface-surface intersections, or poor placement of the gray or white matter surface for any reason.

Of the 934 participants with raw T1-weighted MRI data, 29 participants’ data failed the QC procedures. Of these 29, 13 participants’ data were excluded before any processing due to severe motion and slicing artifacts. The remaining 16 participants failed the CIVET pipeline for reasons including the presence of bright blood vessels and poor contrast. Thus, 905 participants passed the QC procedures.

### Sample sizes

Of the 526 participants with polygenic score data, 391 had T1-weighted neuroimaging data and 518 had neurocognitive measure data. Thus, 391 and 518 children and adolescents were included in the analyses of CT/SA and neurocognitive skills, respectively.

### Neurocognitive Measures

Participants completed tasks from the NIH Toolbox Cognition Battery including the Flanker Inhibitory Control and Attention (Zelazo et al., 2013), List Sorting Working Memory (Tulsky et al., 2013), Picture Sequence Memory (Bauer et al., 2013; Dikmen et al., 2014), and Picture Vocabulary Tests (Gershon et al., 2013, 2014) (see Supplemental Materials). In addition, parents reported on children’s attention problems. Attention problems were measured using two questionnaire items: parent report of a previous child diagnosis of ADHD and/or parent report of significant child attention problems.

### Statistical Analyses

We first investigated whether PGS-EA and parental education were associated with the outcome of interest (e.g., total SA) in separate models. Then, if they were both significantly associated with the outcome of interest, we examined their unique contributions in regression models in which they were both included as predictors.

To examine associations with global measures of SA (total SA) and CT (mean CT) and neurocognitive measures, multiple linear regression analyses in SAS (version 9.4) were conducted using the general linear model (GLM) procedure. Effect sizes (partial eta squared [*η_p_^2^*]) are presented, with values of .01, .06, and .14 indicating small, medium, and large effects, respectively (Cohen, 1988; Richardson, 2011). Given that interactions between socioeconomic factors and PGS-EA were not significant, they were not included in the final regression models. Logistic regression was used to examine associations of PGS-EA and parental education with attention problems.

Covariates included in the regression models were age, age^2^, gender, and scanner/site. In the PING study, 12 MRI scanners were used across the nine data collection sites. Thus, analyses predicting cortical structure included scanner as a covariate, and analyses predicting neurocognitive skills included site as a covariate. In addition, to minimize the chance of population structure explaining the polygenic score results, we extracted 10 first principal components (PC10) and used them as covariates. Without controlling for those principal components, random differences in population genomic signature can explain outcomes, if different populations also happen to differ in the outcome (Price et al., 2006).

Vertex-level neuroanatomical variables of interest included surface area and thickness at each of 81,924 cortical vertices. To examine associations with PGS-EA (or parental education), general linear models were conducted for each vertex for SA and CT using the SurfStat toolbox (http://www.math.mcgill.ca/keith/surfstat/). At every cortical point, the *t*-statistic for the association between brain structure (SA, CT) and PGS-EA (or parental education) was mapped onto a standard cortical surface. Correction for multiple comparisons using random field theory (RFT) (Worsley et al., 2004) was then applied to the resultant map to determine the regions of cortex significantly associated with PGS-EA (or parental education). To identify significant clusters, an initial height threshold of *p* < .001 was implemented at the vertex level, and a corrected family-wise error (*p* < .05) was then applied. In addition, vertex-level RFT thresholding was implemented using the vertex-wise RFT critical t-value (Worsley et al., 2004). Results focus on brain areas significant at the cluster level and include mention of brain areas significant at the vertex level. Vertex level correction is more stringent than cluster-based correction (Woo et al., 2014).

In addition to the covariates mentioned above, the main vertex-wise analyses also controlled for total brain volume. Supplemental analyses not adjusting for total brain volume are also presented based on current recommendations (Mills et al., 2016; Vijayakumar et al., 2018) and to compare our results with those of previous studies that did not control for global measures (McDermott et al., 2019; Noble et al., 2015).

## Results

### Descriptive Statistics

Descriptive statistics and zero-order correlations are presented in Table 1. PGS-EA was significantly correlated with parental education (*r* = .21, *p* < .0001) and family income (*r* = .10, *p* = .03). PGS-EA data were normally distributed.

**Table 1.** Descriptive statistics and zero-order correlations.

|  | 1 | 2 | 3 | 4 | 5 | 6 | 7 | 8 | 9 | 10 |
| --- | --- | --- | --- | --- | --- | --- | --- | --- | --- | --- |
| 1 PGS-EA | - |  |  |  |  |  |  |  |  |  |
| Parental<br>education |  |  |  |  |  |  |  |  |  |  |
| 2 (years) | .21*** | - |  |  |  |  |  |  |  |  |
| Total SA |  |  |  |  |  |  |  |  |  |  |
| 3 (mm<sup>2</sup>) | .09+ | .05 | - |  |  |  |  |  |  |  |
| Global<br>(average) |  |  |  |  |  |  |  |  |  |  |
| 4 CT (mm) | .04 | .10+ | .22*** | - |  |  |  |  |  |  |
| 5 Vocabulary | .03 | .03 | .10* | -.52*** | - |  |  |  |  |  |
| Inhibitory |  |  |  |  |  |  |  |  |  |  |
| 6 control | -.03 | -.03 | .17*** | -.48*** | .72*** | - |  |  |  |  |
| Working |  |  |  |  |  |  |  |  |  |  |
| 7 memory | .001 | -.003 | .16** | -.43*** | .77*** | .77*** | - |  |  |  |
| Sustained |  |  |  |  |  |  |  |  |  |  |
| 8 attention | -.02 | -.01 | .18*** | -.46*** | .67*** | .96*** | .71*** | - |  |  |
| Episodic |  |  |  |  |  |  |  |  |  |  |
| 9 memory | .03 | -.01 | .04 | -.49*** | .70*** | .70*** | .76*** | .65*** | - |  |
| Attention |  |  |  |  |  |  |  |  |  |  |
| 10 problems | -.12** | -.04 | .02 | -.005 | -.01 | .04 | .03 | .06 | -.05 | - |
| *N* | 526 | 503 | 391 | 391 | 518 | 513 | 516 | 515 | 519 | 510 |
|  | .000043 | 15.73 | 200523.00 | 3.11 | .89 | 7.63 | 17.95 | 8.08 | 26.22 | 47<sup>a</sup> |
| *M (SD)* | (.000193) | (1.86) | (16564.94) | (.17) | (1.40) | (1.84) | (5.34) | (1.75) | (11.08) | (9.22<sup>b</sup>) |

*Note.* PGS-EA, polygenic score for educational attainment; SA, cortical surface area; CT, cortical thickness. \* $p < .05$ ; \*\* $p < .01$ ; \*\*\* $p < .001$ ; + $p < .10$
<sup>a</sup> Number of children with attention problems
<sup>b</sup> Percentage of children with attention problems

### PGS-EA, Parental Education, and SA

#### PGS-EA

PGS-EA was significantly positively associated with total SA, *β* = .11, *p* = .0083, *η_p_^2^* = .0187, and remained significantly associated with total SA after accounting for parental education, *β* = .09, *p* = .0279, *ηp* = .0134. Vertex-wise analyses indicated that PGS-EA were significantly (*p* < .05, RFT-corrected) associated with SA in the left medial orbitofrontal gyrus and left inferior frontal gyrus (see Figure 1 and Table 2). These associations remained significant after controlling for parental education (see Figure 1 and Table 2). A consistent pattern of results was found when examining PGS-EA computed at different *p*-value thresholds.

**Figure 1.**
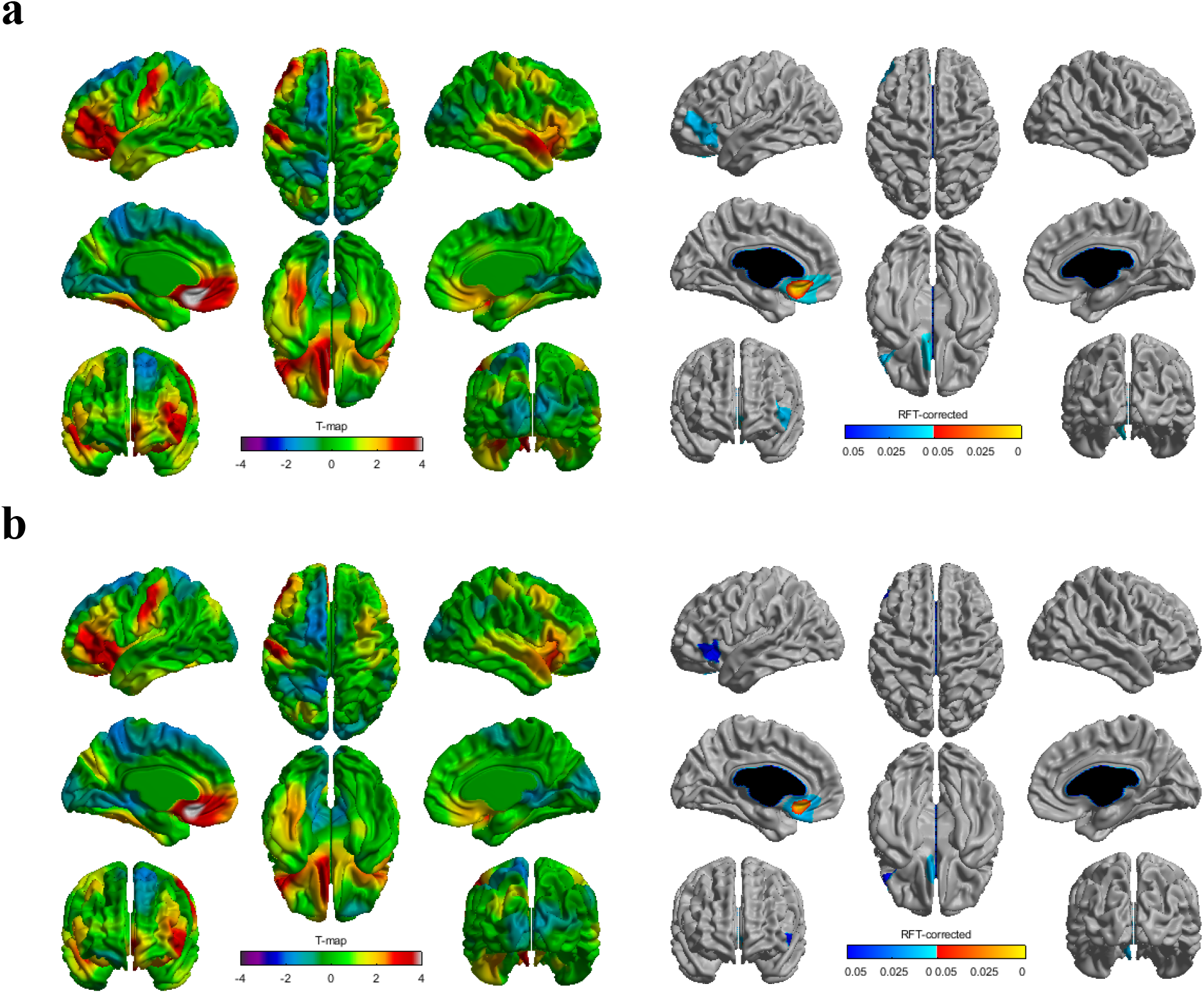
Higher polygenic scores for educational attainment (PGS-EA) were associated with greater cortical surface area (SA) in children and adolescents **(a)** without adjusting for parental education and **(b)** while adjusting for parental education. The left and right panels show *t*-statistics and *p* values (*p* < .05 after correcting for multiple comparisons using random field theory), respectively. Covariates were age, age^2^, gender, scanner, principal components 1-10, and total brain volume. On the right panel, cool colors correspond to areas significant at the cluster level and warm colors correspond to areas significant at the vertex level. RFT, random field theory

**Table 2.**
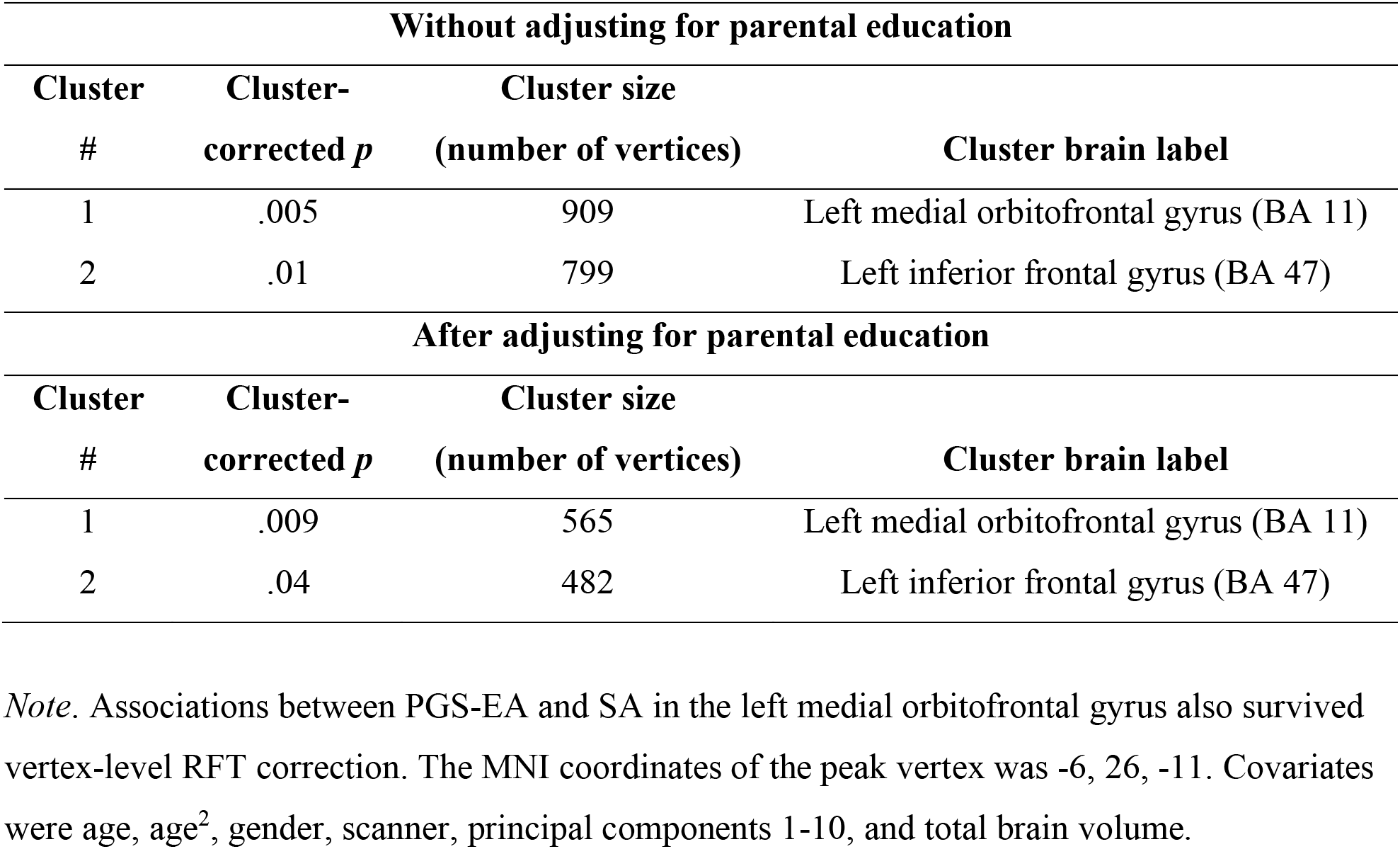
Clusters for significant associations between polygenic scores for educational attainment (PGS-EA) and cortical surface area (SA)

| Without adjusting for parental education |  |  |  |
| --- | --- | --- | --- |
| Cluster # | Cluster-corrected $p$ | Cluster size (number of vertices) | Cluster brain label |
| 1 | .005 | 909 | Left medial orbitofrontal gyrus (BA 11) |
| 2 | .01 | 799 | Left inferior frontal gyrus (BA 47) |
| After adjusting for parental education |  |  |  |
| Cluster # | Cluster-corrected $p$ | Cluster size (number of vertices) | Cluster brain label |
| 1 | .009 | 565 | Left medial orbitofrontal gyrus (BA 11) |
| 2 | .04 | 482 | Left inferior frontal gyrus (BA 47) |

*Note.* Associations between PGS-EA and SA in the left medial orbitofrontal gyrus also survived vertex-level RFT correction. The MNI coordinates of the peak vertex was -6, 26, -11. Covariates were age, age<sup>2</sup>, gender, scanner, principal components 1-10, and total brain volume.

#### Parental education

Parental education was significantly positively associated with total SA, *β* = .13, *p* = .0022, *ηp* = .0257, and remained significantly associated with total SA after accounting for PGS-EA, *β* = .12, *p* = .0076, *η_p_^2^* = .0196. Vertex-wise analyses indicated that higher parental education was significantly (*p* < .05, RFT-corrected) associated with greater SA in the left fusiform gyrus and right superior temporal gyrus (see Figure 2 and Table 3). Parental education was significantly associated with SA in the left parahippocampal gyrus after controlling for PGS-EA (see Figure 2 and Table 3).

**Figure 2.**
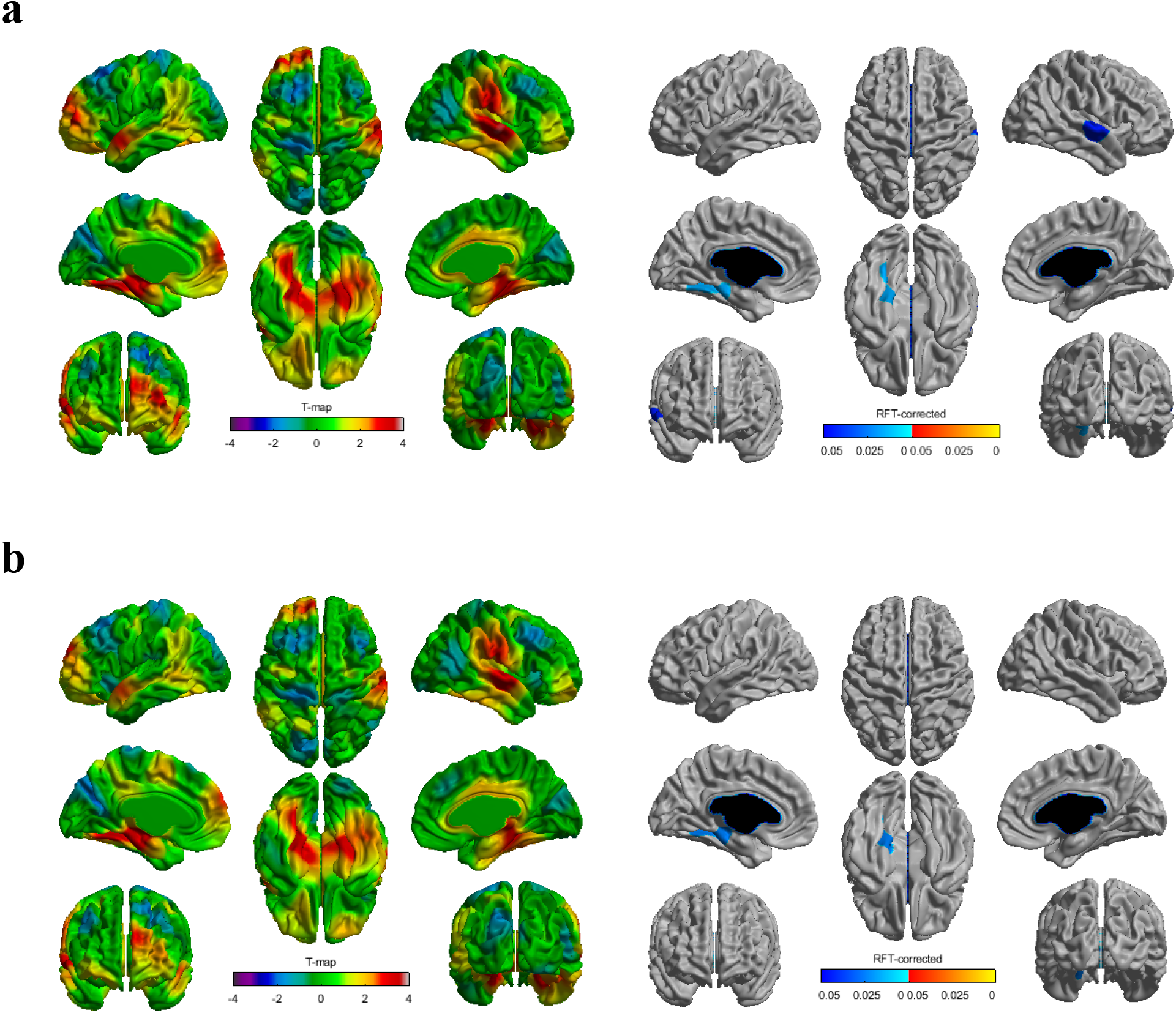
Higher parental education was associated with greater cortical surface area (SA) in children and adolescents **(a)** without adjusting for polygenic score for educational attainment (PGS-EA) and **(b)** while adjusting for PGS-EA. The left and right panels show *t*-statistics and *p* values (*p* < .05 after correcting for multiple comparisons using random field theory), respectively. Covariates were age, age^2^, gender, scanner, and total brain volume. Models including PGS-EA also adjusted for principal components 1-10. On the right panel, cool colors correspond to areas significant at the cluster level and warm colors correspond to areas significant at the vertex level. RFT, random field theory

**Table 3.** Clusters for significant associations between parental education and cortical surface area (SA)

| **Without adjusting for PGS-EA** |  |  |  |
| --- | --- | --- | --- |
| **Cluster #** | **Cluster-corrected *p*** | **Cluster size (number of vertices)** | **Cluster brain label** |
| 1 | .015 | 574 | Left fusiform gyrus (BA 37) |
| 2 | .044 | 493 | Right superior temporal gyrus (BA 21) |
| **After adjusting for PGS-EA** |  |  |  |
| **Cluster #** | **Cluster-corrected *p*** | **Cluster size (number of vertices)** | **Cluster brain label** |
| 1 | .024 | 503 | Left parahippocampal gyrus (BA 36) |

*Note.* Covariates were age, age<sup>2</sup>, gender, scanner, total brain volume, and principal components 1-10.
PGS-EA, polygenic score for educational attainment

### PGS-EA, Parental Education, and CT

PGS-EA was not significantly associated with global CT, *β* = .04, *p* = .2195, or regional CT. Parental education was not significantly associated with global CT, *β* = .05, *p* = .1003, or regional CT.

### PGS-EA, Parental Education, and Neurocognitive Measures

#### PGS-EA

PGS-EA was significantly positively associated with vocabulary, working memory, and episodic memory and negatively associated with attention problems (see Table S2). PGS-EA remained significantly associated with vocabulary, episodic memory, and attention problems after controlling for parental education (see Table S2). A similar pattern of results was found at different *p*-value thresholds with the exception that PGS-EA at other *p*-value thresholds was more strongly associated with sustained attention and working memory (see Table S4).

#### Parental education

Higher parental education was significantly associated with greater inhibitory control, working memory, sustained attention, vocabulary, and episodic memory (see Table S2). Associations with all but episodic memory remained significant after controlling for PGS-EA (see Table S2).

### Supplemental Analyses

A similar pattern of results emerged when not controlling for total brain volume but with significant associations between parental education and SA in more cortical regions (see Figures S1 and S2 and Tables S5 and S6).

## Discussion

The goal of this study was to examine the independent associations of polygenic scores for educational attainment (PGS-EA) and parental education with cortical structure and neurocognitive measures in children and adolescents. Replicating previous findings (Belsky et al., 2016, 2018; Judd et al., 2020; Selzam et al., 2017), higher parental education was significantly correlated with higher PGS-EA in children and adolescents, which may indicate a gene-environment correlation. For example, passive gene-environment correlation may occur because parents create a family environment that corresponds to their genotypes and correlates with the genotypes of their children. These associations underscore the importance of accounting for parental education when analyzing PGS-EA and vice versa. Results from our study, which took this approach, indicated that PGS-EA and parental education explained unique variability in cortical surface area (SA), vocabulary, and attention outcomes.

### PGS-EA and Parental Education are Independently Associated with SA

#### PGS-EA

Higher genome-wide genetic predisposition to educational attainment was significantly associated with greater total SA in children and adolescents, consistent with previous research on adolescents and adults (Judd et al., 2020; Mitchell et al., 2020). Associations between PGS-EA and total SA were attenuated but remained significant after accounting for parental education. These findings suggest that associations between PGS-EA and total SA could arise, in part, from correlations between participants’ PGS-EA and environments associated with their socioeconomic background.

Vertex-wise analyses revealed that PGS-EA were significantly associated with SA in left medial orbitofrontal and inferior frontal regions, and these associations were attenuated but remained significant after additionally controlling for parental education. A previous study of adults (Mitchell et al., 2020) also found PGS-EA to be significantly associated with SA in these frontal regions. Both cognitive and “non-cognitive” skills (e.g., motivation, persistence, grit) are necessary for academic success (Heckman, 2006). PGS-EA have been significantly associated with cognitive (Allegrini et al., 2019; Plomin & von Stumm, 2018; Wertz et al., 2018) and non-cognitive skills (Belsky et al., 2016; Smith-Woolley et al., 2019). The medial orbitofrontal cortex (OFC) has been associated with inhibitory control in emotionally- or motivationally-salient contexts (Rolls, 2019). Genotypes linked to higher educational attainment may impact SA in the medial OFC, leading to greater impulse control and/or academic motivation and in turn school performance. Associations between PGS-EA and SA in the left inferior frontal gyrus, which has been associated with language skills (Friederici, 2011), are consistent with the robust associations found between PGS-EA and language skills (Loughnan et al., 2021; von Stumm et al., 2020).

Genetic variants associated with educational attainment have been linked with genes showing elevated expression in neural tissue (Okbay et al., 2016). Genetic propensity to higher educational attainment may include variants that promote optimal cortical development. The cellular processes underlying developmental changes in SA, including synaptic function, have been associated with genes linked with the significant SNPs identified in GWAS of educational attainment (Deary et al., 2021; Okbay et al., 2016).

#### Parental education

Higher parental education was significantly associated with greater total SA, replicating previous findings using the PING dataset (Noble et al., 2015) and other large-scale datasets (Judd et al., 2020; McDermott et al., 2019). Extending previous work, our findings indicated that this association was attenuated after adjusting for PGS-EA, suggesting genetic confounding (Wertz et al., 2020). The fact that parental education remained significantly associated with total SA after accounting for PGS-EA leaves open the possibility of environmental transmission.

Vertex-wise analyses indicated that associations between parental education and SA were most prominent in the left parahippocampal/fusiform gyrus and right superior temporal gyrus. Parental education was significantly associated with SA in the left parahippocampal gyrus after adjusting for PGS-EA. The parahippocampal gyrus, as part of the medial temporal lobe, has been strongly associated with episodic memory (Eichenbaum, 2006), which varies significantly across socioeconomic gradients (Noble et al., 2005, 2007; Noble & Giebler, 2020). When not adjusting for total brain volume, similar to analytic approaches used in previous work (McDermott et al., 2019; Noble et al., 2015), parental education was significantly associated with SA in more cortical regions, including larger portions of the bilateral parahippocampal gyrus, left fusiform gyrus, and right superior temporal gyrus (see Figure S2).

These findings are consistent with the notion that socioeconomic differences in SA are partially driven by environmental transmission, although inferences about environmental transmission cannot be made. Socioeconomic factors may impact SA in children and adolescents via multiple proximal environmental factors. For example, socioeconomic disadvantage has been consistently associated with exposure to chronic stressors (e.g., household chaos and unpredictability, neighborhood violence, crowding/noise, family conflict) and reduced quantity and quality of language input to children (Duncan et al., 2017; Evans & Kim, 2013; Merz et al., 2019; Pace et al., 2017). Evidence from randomized trials of poverty reduction and animal models of chronic stress and environmental enrichment suggests that at least part of the association between socioeconomic factors and children’s cognitive development may be environmentally mediated (Davidson & McEwen, 2012; Duncan et al., 2017; van Praag et al., 2000). In the current study, associations between parental education and SA in children and adolescents, even after accounting for PGS-EA, cannot be interpreted as environmental effects. Other sources of genetic confounding may play a role in those associations (Wertz et al., 2020).

PGS-EA and parental education were not significantly associated with global or regional CT, consistent with a previous study of adolescents (Judd et al., 2020). Similarly, findings linking family SES with CT have been variable, with some studies indicating significant correlations, but others reporting no links (Noble & Giebler, 2020). These findings may suggest differential associations of parental education and PGS-EA with SA and CT, which would suggest that these associations may rely more heavily on certain underlying cellular mechanisms than others (Panizzon et al., 2009; Rakic, 1988; Raznahan et al., 2011).

### PGS-EA and Parental Education are Independently Associated with Neurocognitive Measures

#### PGS-EA

Associations of PGS-EA with vocabulary, episodic memory, and reduced attention problems were attenuated but remained significant after controlling for parental education. Associations between educational attainment polygenic scores and these neurocognitive skills may partially explain associations of PGS-EA with general cognitive ability and academic achievement (von Stumm et al., 2020). Results linking higher PGS-EA with lower risk for attention problems are consistent with associations between PGS-EA and SA in the medial OFC. In addition, associations between higher PGS-EA and higher vocabulary are consistent with links between PGS-EA and SA in the left inferior frontal gyrus (Friederici, 2011).

#### Parental education

Higher parental education was significantly associated with higher inhibitory control, working memory, sustained attention, and vocabulary. These associations were attenuated after accounting for genetic predisposition to educational attainment. These results are consistent with prior work indicating socioeconomic disparities in language and executive function in children (Lawson et al., 2017; Noble et al., 2005, 2007). The current findings extend this work by showing that such associations remain significant even after controlling for children’s education-linked genetics. These results are consistent with the notion that SES-related environmental factors (e.g., language input) may be associated with language and executive function above and beyond genetic factors (Duncan et al., 2017).

Several limitations of this study must be taken into account when interpreting the findings. First, due to the cross-sectional, correlational design of this study, causal inferences cannot be made. Second, as in most studies that use polygenic scores (Elliott et al., 2019; von Stumm et al., 2020), analyses included only participants of European ancestry. Large-scale GWAS, which are required for identifying genetic variants that are reliably associated with a phenotype, are currently not available in populations with other ancestries. Thus, findings from this study are not generalizable to other ethnicities.

Findings from this study indicated that education-associated genetics and parental education accounted for unique variance in SA in children and adolescents. PGS-EA and parental education were most prominently associated with SA in frontal and temporal regions. These results shed light on the role of education-linked genetics in contributing to brain structure and neurocognitive skills in children and adolescents.

## Acknowledgements

Data used in this study are available through the National Institute of Mental Health (NIMH) Data Archive (https://nda.nih.gov/) after providing the required data user agreement.

## Supplemental Materials

### NIH Toolbox Cognition Battery Tasks

#### Vocabulary

During the Picture Vocabulary Test (Gershon et al., 2013, 2014), participants listened to an audio recording of a word on each trial, while viewing four images on a computer screen. Participants were asked to select the image on the computer screen that best matched the meaning of the spoken word. Through computer adaptive testing, the difficulty of the words presented was tailored to the ability of the participant. Test difficulty is adapted such that participants have a 50% chance of answering correctly on each trial.

#### Working memory

On the List Sorting Working Memory test (Tulsky et al., 2013), a series of pictures of different foods and animals were presented on a computer screen visually and aurally, one at a time. In the one-list condition, participants were told to remember stimuli from one category (food or animals) and repeat them in size order, from smallest to largest. In the two-list condition, participants were told to remember stimuli from two categories (food and animals, intermixed) and then report the food in size order, followed by the animals in size order. Working memory scores consisted of the total items correct across the one- and two-list conditions.

#### Inhibitory control

On the Flanker Inhibitory Control and Attention test (Zelazo et al., 2013), participants were asked to focus on the central stimulus while inhibiting attention to the flanker (surrounding) stimuli. The test consisted of a block of 25 fish trials followed by a block of 25 arrow trials, with 16 congruent and 9 incongruent trials in each block, presented in pseudorandom order. On congruent trials, all the stimuli were pointing in the same direction (right or left). On incongruent trials, the central stimulus was pointing in the opposite direction of the flanker stimuli. Congruent and incongruent trials were intermixed in each block of test trials. Performance on both congruent and incongruent trials was recorded. A two-vector scoring method was used that incorporated both accuracy and reaction time for participants who maintained a high level of accuracy (>80% correct), and accuracy only for those who did not meet this criterion.

#### Attention

Attention was measured through participants’ performance on the Flanker Inhibitory Control and Attention test (Zelazo et al., 2013) and parent-reported attention problems. Performance on congruent trials of the Flanker task was used as an index of sustained attention (Akshoomoff et al., 2014).

#### Episodic memory

On the Picture Sequence Memory Test (Bauer et al., 2013; Dikmen et al., 2014), participants viewed a sequence of thematically related pictures that appeared one at a time in the center of the computer screen (2 s each). As each picture appeared, an audio recording described the content of the picture. After each picture was presented, it was moved to a unique spatial position on the computer screen that matched the temporal order in which the pictures were presented. After all the pictures had been presented and moved to their unique spatial position, they disappeared from the screen. Three seconds later, all the pictures re-appeared on the computer screen in a scrambled order, and participants were asked to move each picture back to its correct spatial position. Picture sequence length varied from 6 to 15 pictures depending on the age of the participant.

### Results for Family Income

#### Family Income, PGS-EA, and SA

Higher family income was significantly associated with greater total SA, *β* = .10, *p* = .0173, *η_p_^2^ =* .0156. This association remained significant after controlling for PGS-EA, *β* = .09, *p* = .0306, *η_p_^2^* = .0129. Vertex-wise analyses revealed that higher family income was significantly (*p* < .05, RFT-corrected) associated with greater SA in the right inferior temporal gyrus and fusiform gyrus.

#### Family Income, PGS-EA, and CT

Family income was significantly positively associated with average CT, *β* = .07, *p* = .0183, *η_p_^2^* = .0149. Vertex-wise analyses indicated that family income was not associated with CT in any specific cortical regions.

#### Family Income, PGS-EA, and Neurocognitive Measures

Higher family income was significantly associated with higher inhibitory control, working memory, sustained attention, vocabulary, and episodic memory (see Table S3). Higher family income was significantly associated with inhibitory control, sustained attention, and vocabulary after controlling for PGS-EA (see Table S3).

---

**Table S1.** The number of SNPs from the Lee et al. (2018) (EA3) GWAS included in each PGS-EA *p*-value threshold.

| **$p$-value threshold** | **Number of SNPs from Lee et al. (2018) included** |
| --- | --- |
| $p < 5 \times 10^{-8}$ | 604 |
| $p < 1 \times 10^{-7}$ | 694 |
| $p < 1 \times 10^{-6}$ | 1067 |
| $p < 1 \times 10^{-5}$ | 1795 |
| $p < .0001$ | 3241 |
| $p < .001$ | 6395 |
| $p < .01$ | 14426 |
| $p < .05$ | 28535 |
| $p < .1$ | 38875 |
| $p < .5$ | 80125 |
| $p < 1$ | 99493 |

**Table S2.**
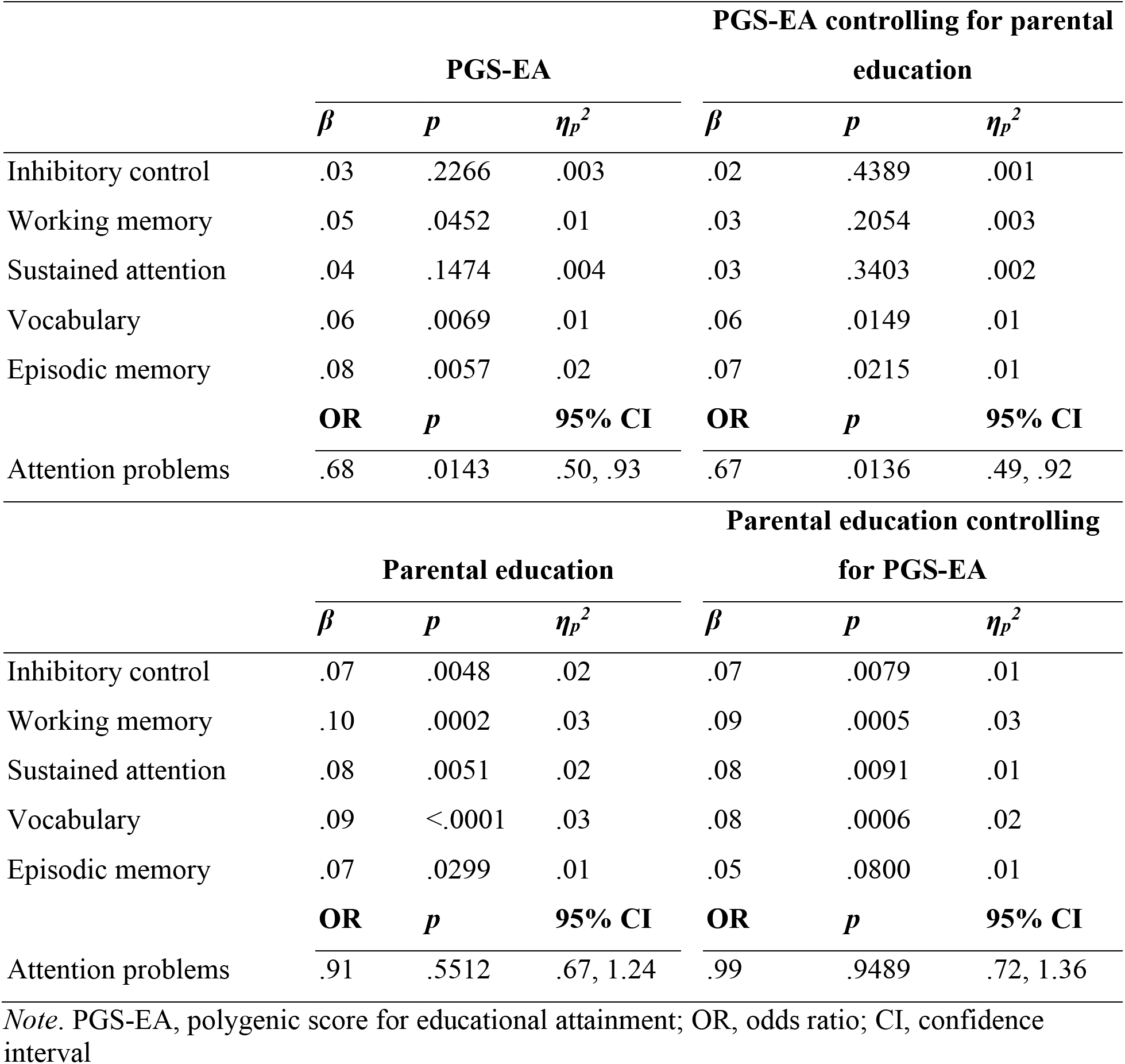
Associations of PGS-EA and parental education with neurocognitive measures.

**Table S3.** Associations between family income and neurocognitive measures.

|  | Family income |  |  | Family income controlling for PGS-EA |  |  |
| --- | --- | --- | --- | --- | --- | --- |
| | $\beta$ | $p$ | $\eta_p^2$ | $\beta$ | $p$ | $\eta_p^2$ |
| Inhibitory control | .06 | .0143 | .01 | .06 | .0184 | .01 |
| Working memory | .05 | .0468 | .01 | .05 | .0687 | .01 |
| Sustained attention | .06 | .0337 | .01 | .06 | .0442 | .01 |
| Vocabulary | .06 | .0069 | .02 | .06 | .0118 | .01 |
| Episodic memory | .06 | .0389 | .01 | .05 | .0637 | .01 |
|  | **OR** | **$p$** | **95% CI** | **OR** | **$p$** | **95% CI** |
| Attention problems | .92 | .60 | .69, 1.24 | .96 | .77 | .71, 1.29 |

*Note.* PGS-EA, polygenic score for educational attainment; OR, odds ratio; CI, confidence interval

**Table S4.** Associations of PGS-EA computed at different *p*-value thresholds with neurocognitive skills.

| **PGS-EA $p$-value threshold** | **Neurocognitive skill** | **$\beta$** | **$p$** |
| --- | --- | --- | --- |
| **$p < 5 \times 10^{-8}$** | Inhibitory control | .05 | .0329 |
|  | Working memory | .10 | .0003 |
|  | Sustained attention | .08 | .0043 |
|  | Vocabulary | .14 | <.0001 |
|  | Episodic memory | .13 | <.0001 |
| **$p < 1 \times 10^{-6}$** | Inhibitory control | .04 | .1354 |
|  | Working memory | .06 | .0116 |
|  | Sustained attention | .05 | .1128 |
|  | Vocabulary | .07 | .0009 |
|  | Episodic memory | .11 | .0002 |
| **$p < 1 \times 10^{-5}$** | Inhibitory control | .04 | .0944 |
|  | Working memory | .08 | .0028 |
|  | Sustained attention | .04 | .1393 |
|  | Vocabulary | .09 | .0001 |
|  | Episodic memory | .10 | .0003 |
| **$p < .0001$** | Inhibitory control | .06 | .0240 |
|  | Working memory | .08 | .0027 |
|  | Sustained attention | .06 | .0281 |
|  | Vocabulary | .10 | <.0001 |
|  | Episodic memory | .12 | <.0001 |
| **$p < .001$** | Inhibitory control | .06 | .0248 |
|  | Working memory | .08 | .0022 |
|  | Sustained attention | .06 | .0282 |
|  | Vocabulary | .11 | <.0001 |
|  | Episodic memory | .12 | <.0001 |
| **$p < .01$** | Inhibitory control | .05 | .0417 |
|  | Working memory | .08 | .0013 |
|  | Sustained attention | .07 | .0243 |
|  | Vocabulary | .12 | <.0001 |
|  | Episodic memory | .13 | <.0001 |
| **$p < .05$** | Inhibitory control | .05 | .0512 |
|  | Working memory | .10 | .0002 |
|  | Sustained attention | .07 | .0275 |
|  | Vocabulary | .13 | <.0001 |
|  | Episodic memory | .13 | <.0001 |
| **$p < .1$** | Inhibitory control | .05 | .0360 |
|  | Working memory | .09 | .0007 |
|  | Sustained attention | .08 | .0095 |
|  | Vocabulary | .13 | <.0001 |
|  | Episodic memory | .12 | .0001 |
| **$p < 1$** | Inhibitory control | .05 | .0342 |
|  | Working memory | .10 | .0003 |
|  | Sustained attention | .08 | .0048 |
|  | Vocabulary | .14 | <.0001 |
|  | Episodic memory | .13 | <.0001 |

*Note.* Results for $p < 1 \times 10^{-7}$ are presented in the main manuscript file. Covariates were age, age<sup>2</sup>, gender, site, and principal components 1-10.

**Table S5.**
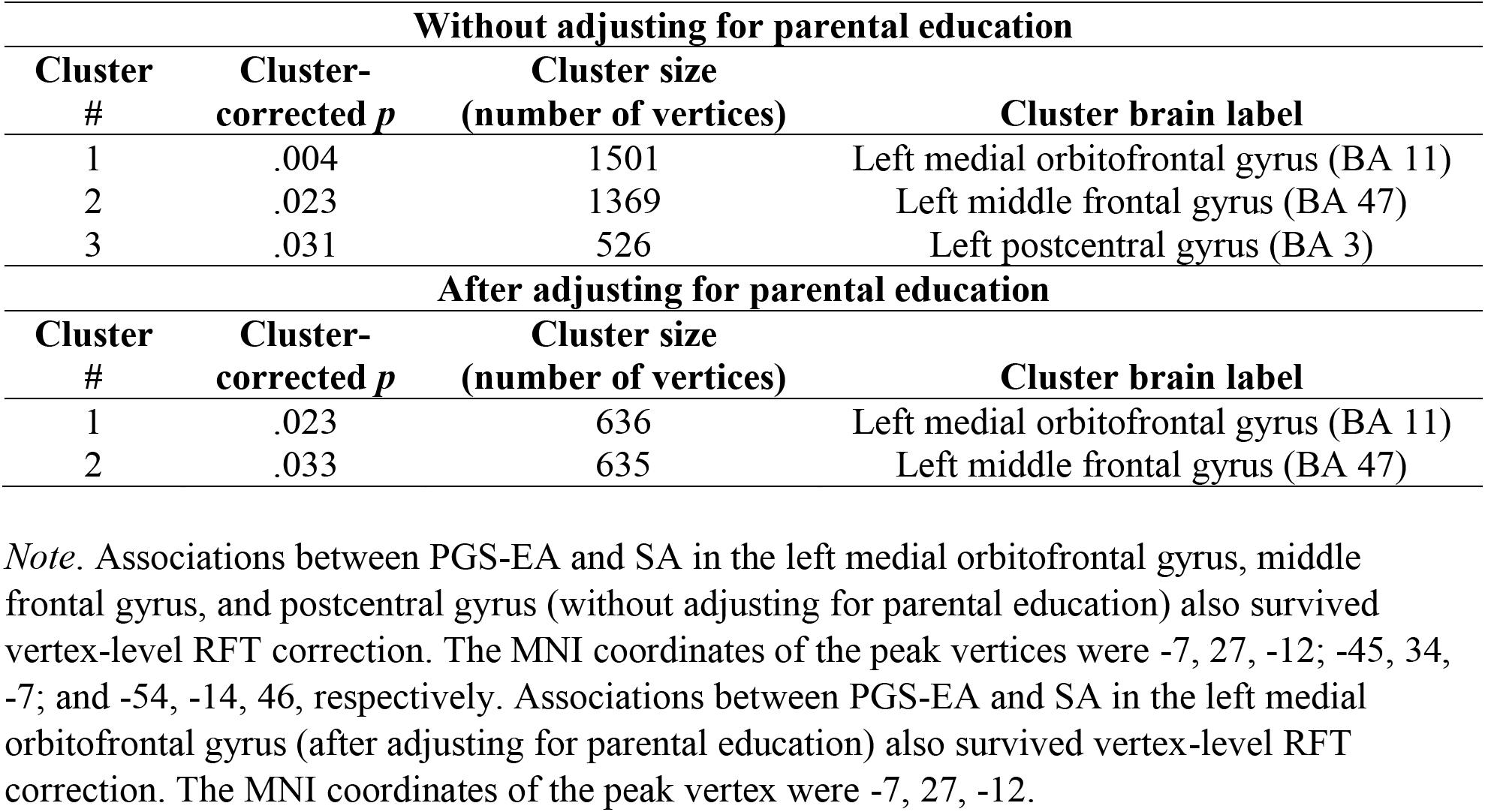
Clusters for significant associations between polygenic scores for educational attainment (PGS-EA) and cortical surface area (SA)

| **Without adjusting for parental education** |  |  |  |
| --- | --- | --- | --- |
| **Cluster #** | **Cluster-corrected $p$** | **Cluster size (number of vertices)** | **Cluster brain label** |
| 1 | .004 | 1501 | Left medial orbitofrontal gyrus (BA 11) |
| 2 | .023 | 1369 | Left middle frontal gyrus (BA 47) |
| 3 | .031 | 526 | Left postcentral gyrus (BA 3) |
| **After adjusting for parental education** |  |  |  |
| **Cluster #** | **Cluster-corrected $p$** | **Cluster size (number of vertices)** | **Cluster brain label** |
| 1 | .023 | 636 | Left medial orbitofrontal gyrus (BA 11) |
| 2 | .033 | 635 | Left middle frontal gyrus (BA 47) |

*Note.* Associations between PGS-EA and SA in the left medial orbitofrontal gyrus, middle frontal gyrus, and postcentral gyrus (without adjusting for parental education) also survived vertex-level RFT correction. The MNI coordinates of the peak vertices were -7, 27, -12; -45, 34, -7; and -54, -14, 46, respectively. Associations between PGS-EA and SA in the left medial orbitofrontal gyrus (after adjusting for parental education) also survived vertex-level RFT correction. The MNI coordinates of the peak vertex were -7, 27, -12.

**Table S6.** Clusters for significant associations between parental education and cortical surface area (SA)

| **Without adjusting for PGS-EA** |  |  |  |
| --- | --- | --- | --- |
| **Cluster #** | **Cluster-corrected *p*** | **Cluster size (number of vertices)** | **Cluster brain label** |
| 1 | .003 | 1612 | Left parahippocampal, fusiform and lingual |
| 2 | .003 | 3116 | Right superior temporal, supramarginal, postcentral |
| 3 | .008 | 1592 | Right parahippocampal, fusiform |
| 4 | .003 | 953 | Left superior frontal gyrus (BA 10) |
| **After adjusting for PGS-EA** |  |  |  |
| **Cluster #** | **Cluster-corrected *p*** | **Cluster size (number of vertices)** | **Cluster brain label** |
| 1 | .011 | 1277 | Left parahippocampal, fusiform and lingual |
| 2 | .026 | 852 | Right superior temporal |
| 3 | .046 | 948 | Right parahippocampal |
| 4 | .034 | 683 | Right postcentral gyrus (BA 3) |

*Note.* Associations of parental education with the first three clusters (without adjusting for PGS-EA) also survived vertex-level RFT correction. The MNI coordinates of the peak vertices were -30, -47, -10; 63, -6, -3; and 29, -26, -21, respectively. Associations of parental education with the first three clusters (after adjusting for PGS-EA) also survived vertex-level RFT correction. The MNI coordinates of the peak vertices were -32, -40, -7; 63, -9, -3; and 28, -29, -18, respectively.

**Figure S1.**
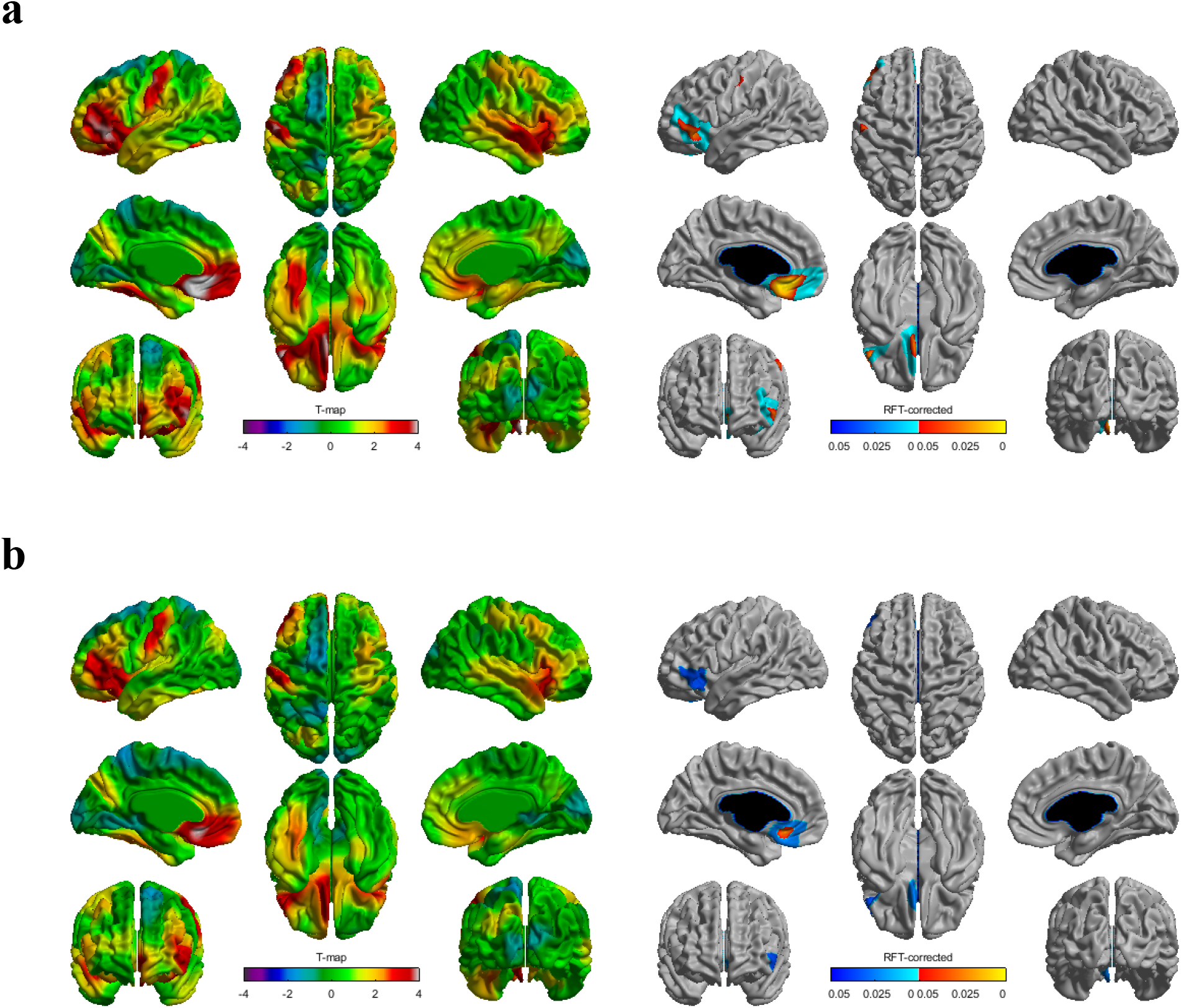
Higher polygenic scores for educational attainment (PGS-EA) were associated with greater cortical surface area (SA) in children and adolescents **(a)** without adjusting for parental education and **(b)** while adjusting for parental education. The left and right panels show *t*-statistics and *p* values (*p* < .05 after correcting for multiple comparisons using random field theory), respectively. Covariates were age, age^2^, gender, scanner, and principal components 1-10.

**Figure S2.**
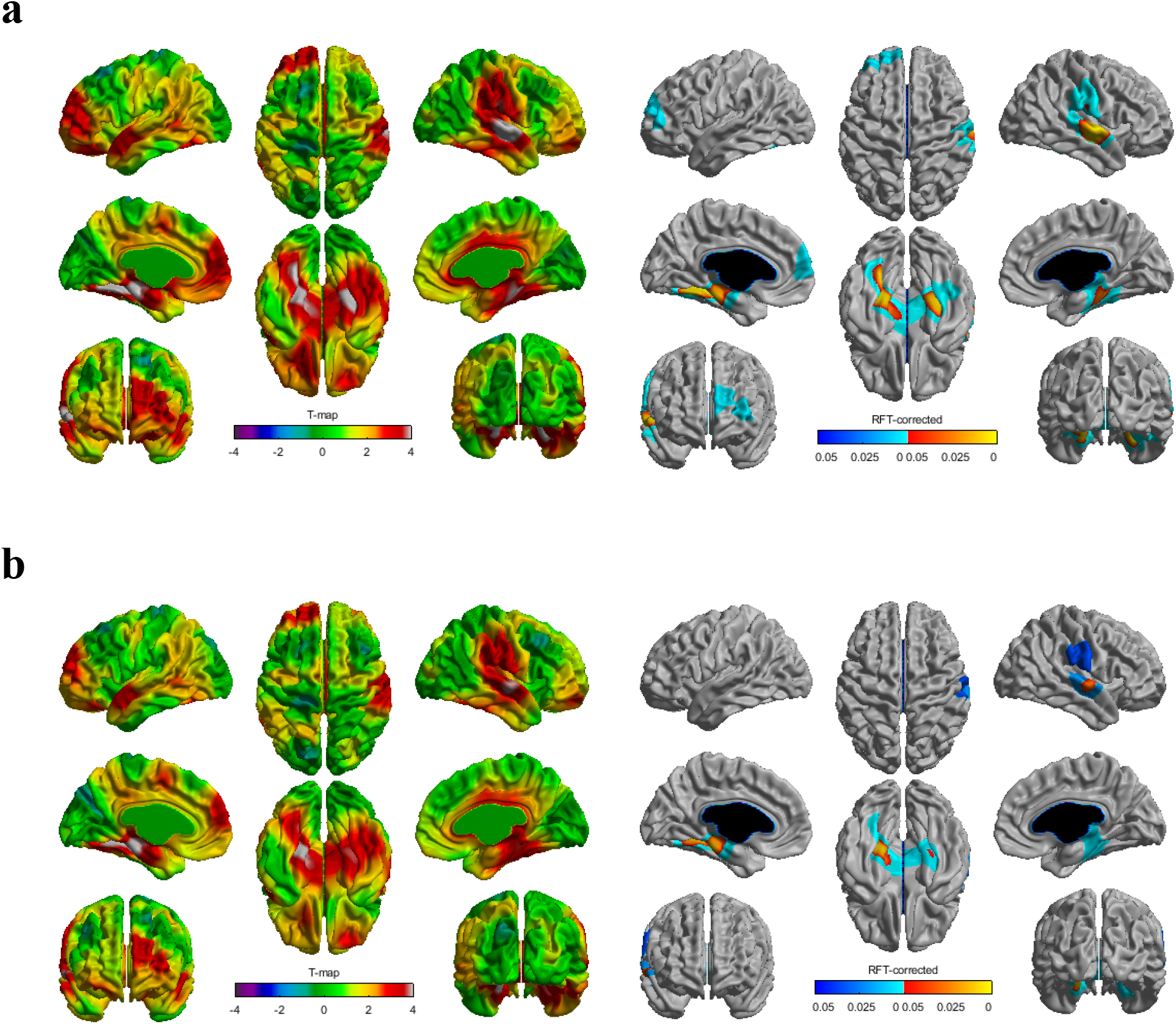
Higher parental education was associated with greater cortical surface area (SA) in children and adolescents **(a)** without adjusting for polygenic score for educational attainment (PGS-EA) and **(b)** while adjusting for PGS-EA. The left and right panels show *t*-statistics and *p* values (*p* < .05 after correcting for multiple comparisons using random field theory), respectively. Covariates were age, age^2^, gender, scanner, and principal components 1-10.

## References

Alemany, S., Jansen, P. R., Muetzel, R. L., Marques, N., El Marroun, H., Jaddoe, V. W. V., Polderman, T. J. C., Tiemeier, H., Posthuma, D., & White, T. (2019). Common Polygenic Variations for Psychiatric Disorders and Cognition in Relation to Brain Morphology in the General Pediatric Population. Journal of the American Academy of Child and Adolescent Psychiatry, 58(6), 600–607. https://doi.org/10.1016/j.jaac.2018.09.443

Allegrini, A. G., Selzam, S., Rimfeld, K., von Stumm, S., Pingault, J. B., & Plomin, R. (2019). Genomic prediction of cognitive traits in childhood and adolescence. Molecular Psychiatry, 24(6), 819–827. https://doi.org/10.1038/s41380-019-0394-4

Armstrong-Carter, E., Wertz, J., & Domingue, B. W. (2021). Genetics and Child Development: Recent Advances and Their Implications for Developmental Research. Child Development Perspectives, 15(1), 57–64. https://doi.org/10.1111/cdep.12400

Bauer, P. J., Dikmen, S. S., Heaton, R. K., Mungas, D., Slotkin, J., & Beaumont, J. L. (2013). III. NIH TOOLBOX COGNITION BATTERY (CB): MEASURING EPISODIC MEMORY. Monographs of the Society for Research in Child Development, 78(4), 34–48. https://doi.org/10.1111/mono.12033

Belsky, D. W., Domingue, B. W., Wedow, R., Arseneault, L., Boardman, J. D., Caspi, A., Conley, D., Fletcher, J. M., Freese, J., Herd, P., Moffitt, T. E., Poulton, R., Sicinski, K., Wertz, J., & Harris, K. M. (2018). Genetic analysis of social-class mobility in five longitudinal studies. Proceedings of the National Academy of Sciences of the United States of America, 115(31), E7275–E7284. https://doi.org/10.1073/pnas.1801238115

Belsky, D. W., Moffitt, T. E., Corcoran, D. L., Domingue, B., Harrington, H., Hogan, S., Houts, R., Ramrakha, S., Sugden, K., Williams, B. S., Poulton, R., & Caspi, A. (2016). The Genetics of Success: How Single-Nucleotide Polymorphisms Associated With Educational Attainment Relate to Life-Course Development. Psychological Science, 27(7), 957–972. https://doi.org/10.1177/0956797616643070

Chang, C. C., Chow, C. C., Tellier, L. C., Vattikuti, S., Purcell, S. M., & Lee, J. J. (2015). Second-generation PLINK: Rising to the challenge of larger and richer datasets. GigaScience, 4, 7. https://doi.org/10.1186/s13742-015-0047-8

Cohen, J. (1988). Statistical Power Analysis for the Behavioral Sciences (2nd ed.). Routledge. https://doi.org/10.4324/9780203771587

Davidson, R. J., & McEwen, B. S. (2012). Social influences on neuroplasticity: Stress and interventions to promote well-being. Nature Neuroscience, 15(5), 689–695. https://doi.org/10.1038/nn.3093

Deary, I. J., Cox, S. R., & Hill, W. D. (2021). Genetic variation, brain, and intelligence differences. Molecular Psychiatry. https://doi.org/10.1038/s41380-021-01027-y

Dikmen, S. S., Bauer, P. J., Weintraub, S., Mungas, D., Slotkin, J., Beaumont, J. L., Gershon, R., Temkin, N. R., & Heaton, R. K. (2014). Measuring Episodic Memory Across the Lifespan: NIH Toolbox Picture Sequence Memory Test. Journal of the International Neuropsychological Society, 20(6), 611–619. https://doi.org/10.1017/S1355617714000460

Domingue, B. W., Belsky, D., Conley, D., Harris, K. M., & Boardman, J. D. (2015). Polygenic Influence on Educational Attainment: New evidence from The National Longitudinal Study of Adolescent to Adult Health. AERA Open, 1(3), 1–13. https://doi.org/10.1177/2332858415599972

Du Rietz, E., Coleman, J., Glanville, K., Choi, S. W., O’Reilly, P. F., & Kuntsi, J. (2018). Association of Polygenic Risk for Attention-Deficit/Hyperactivity Disorder With Co-occurring Traits and Disorders. Biological Psychiatry. Cognitive Neuroscience and Neuroimaging, 3(7), 635–643. https://doi.org/10.1016/j.bpsc.2017.11.013

Duncan, G. J., & Magnuson, K. (2012). Socioeconomic status and cognitive functioning: Moving from correlation to causation. Wiley Interdisciplinary Reviews: Cognitive Science, 3(3), 377–386. https://doi.org/10.1002/wcs.1176

Duncan, G. J., Magnuson, K., & Votruba-Drzal, E. (2017). Moving Beyond Correlations in Assessing the Consequences of Poverty. Annual Review of Psychology, 68, 413–434. https://doi.org/10.1146/annurev-psych-010416-044224

Eichenbaum, H. (2006). Remembering: Functional Organization of the Declarative Memory System. Current Biology, 16(16), R643–R645. https://doi.org/10.1016/j.cub.2006.07.026

Elliott, M. L., Belsky, D. W., Anderson, K., Corcoran, D. L., Ge, T., Knodt, A., Prinz, J. A., Sugden, K., Williams, B., Ireland, D., Poulton, R., Caspi, A., Holmes, A., Moffitt, T., & Hariri, A. R. (2019). A Polygenic Score for Higher Educational Attainment is Associated with Larger Brains. Cerebral Cortex (New York, NY), 29(8), 3496–3504. https://doi.org/10.1093/cercor/bhy219

Euesden, J., Lewis, C. M., & O’Reilly, P. F. (2015). PRSice: Polygenic Risk Score software. Bioinformatics (Oxford, England), 31(9), 1466–1468. https://doi.org/10.1093/bioinformatics/btu848

Evans, G. W., & Kim, P. (2013). Childhood Poverty, Chronic Stress, Self-Regulation, and Coping. Child Development Perspectives, 7(1), 43–48. https://doi.org/10.1111/cdep.12013

Farah, M. J. (2017). The Neuroscience of Socioeconomic Status: Correlates, Causes, and Consequences. Neuron, 96(1), 56–71. https://doi.org/10.1016/j.neuron.2017.08.034

Figlio, D. N., Freese, J., Karbownik, K., & Roth, J. (2017). Socioeconomic status and genetic influences on cognitive development. Proceedings of the National Academy of Sciences of the United States of America, 114(51), 13441–13446. https://doi.org/10.1073/pnas.1708491114

Friederici, A. D. (2011). The brain basis of language processing: From structure to function. Physiological Reviews, 91(4), 1357–1392. https://doi.org/10.1152/physrev.00006.2011

Gershon, R. C., Cook, K. F., Mungas, D., Manly, J. J., Slotkin, J., Beaumont, J. L., & Weintraub, S. (2014). Language Measures of the NIH Toolbox Cognition Battery. Journal of the International Neuropsychological Society, 20(6), 642–651. https://doi.org/10.1017/S1355617714000411

Gershon, R. C., Slotkin, J., Manly, J. J., Blitz, D. L., Beaumont, J. L., Schnipke, D., Wallner-Allen, K., Golinkoff, R. M., Gleason, J. B., Hirsh-Pasek, K., Adams, M. J., & Weintraub, S. (2013). IV. NIH Toolbox Cognition Battery (CB): Measuring Language (Vocabulary Comprehension and Reading Decoding). Monographs of the Society for Research in Child Development, 78(4), 49–69. https://doi.org/10.1111/mono.12034

Heckman, J. J. (2006). Skill formation and the economics of investing in disadvantaged children. Science, 312(5782), 1900–1902. https://doi.org/10.1126/science.1128898

Jernigan, T. L., Brown, T. T., Hagler Jr., D. J., Akshoomoff, N., Bartsch, H., Newman, E., Thompson, W. K., Bloss, C. S., Murray, S. S., Schork, N., Kennedy, D. N., Kuperman, J. M., McCabe, C., Chung, Y., Libiger, O., Maddox, M., Casey, B. J., Chang, L., Ernst, T. M., … Dale, A. M. (2016). The Pediatric Imaging, Neurocognition, and Genetics (PING) Data Repository. NeuroImage, 124, *Part B*, 1149–1154. https://doi.org/10.1016/j.neuroimage.2015.04.057

Judd, N., Sauce, B., Wiedenhoeft, J., Tromp, J., Chaarani, B., Schliep, A., Noort, B. van Penttilä, J., Grimmer, Y., Insensee, C., Becker, A., Banaschewski, T., Bokde, A. L. W., Quinlan, E. B., Desrivières, S., Flor, H., Grigis, A., Gowland, P., Heinz, A., … Klingberg, T. (2020). Cognitive and brain development is independently influenced by socioeconomic status and polygenic scores for educational attainment. Proceedings of the National Academy of Sciences, 117(22), 12411–12418. https://doi.org/10.1073/pnas.2001228117

Khundrakpam, B., Choudhury, S., Vainik, U., Al-Sharif, N., Bhutani, N., Jeon, S., Gold, I., & Evans, A. (2020a). Distinct influence of parental occupation on cortical thickness and surface area in children and adolescents: Relation to self-esteem. Human Brain Mapping, 41(18), 5097–5113. https://doi.org/10.1002/hbm.25169

Khundrakpam, B., Vainik, U., Gong, J., Al-Sharif, N., Bhutani, N., Kiar, G., Zeighami, Y., Kirschner, M., Luo, C., Dagher, A., & Evans, A. (2020b). Neural correlates of polygenic risk score for autism spectrum disorders in general population. Brain Communications, 2(2). https://doi.org/10.1093/braincomms/fcaa092

Krapohl, E., & Plomin, R. (2016). Genetic link between family socioeconomic status and children’s educational achievement estimated from genome-wide SNPs. Molecular Psychiatry, 21(3), 437–443. https://doi.org/10.1038/mp.2015.2

Lawson, G. M., Duda, J. T., Avants, B. B., Wu, J., & Farah, M. J. (2013). Associations between Children’s Socioeconomic Status and Prefrontal Cortical Thickness. Developmental Science, 16(5), 641–652. https://doi.org/10.1111/desc.12096

Lawson, G. M., Hook, C. J., & Farah, M. J. (2017). A meta-analysis of the relationship between socioeconomic status and executive function performance among children. Developmental Science. https://doi.org/10.1111/desc.12529

Lee, J. J., Wedow, R., Okbay, A., Kong, E., Maghzian, O., Zacher, M., Nguyen-Viet, T. A., Bowers, P., Sidorenko, J., Karlsson Linnér, R., Fontana, M. A., Kundu, T., Lee, C., Li, H., Li, R., Royer, R., Timshel, P. N., Walters, R. K., Willoughby, E. A., … Cesarini, D. (2018). Gene discovery and polygenic prediction from a genome-wide association study of educational attainment in 1.1 million individuals. Nature Genetics, 50(8), 1112–1121. https://doi.org/10.1038/s41588-018-0147-3

Loughnan, R. J., Palmer, C. E., Thompson, W. K., Dale, A. M., Jernigan, T. L., & Fan, C. C. (2021). Gene-experience correlation during cognitive development: Evidence from the Adolescent Brain Cognitive Development (ABCD) StudySM. BioRxiv, 637512. https://doi.org/10.1101/637512

Mackey, A. P., Finn, A. S., Leonard, J. A., Jacoby Senghor, D. S., West, M. R., Gabrieli, C. F. O., & Gabrieli, J. D. E. (2015). Neuroanatomical Correlates of the Income Achievement Gap. Psychological Science, 26(6), 925–933. https://doi.org/10.1177/0956797615572233

McCarthy, S., Das, S., Kretzschmar, W., Delaneau, O., Wood, A. R., Teumer, A., Kang, H. M., Fuchsberger, C., Danecek, P., Sharp, K., Luo, Y., Sidore, C., Kwong, A., Timpson, N., Koskinen, S., Vrieze, S., Scott, L. J., Zhang, H., Mahajan, A., … Haplotype Reference Consortium. (2016). A reference panel of 64,976 haplotypes for genotype imputation. Nature Genetics, 48(10), 1279–1283. https://doi.org/10.1038/ng.3643

McDermott, C. L., Seidlitz, J., Nadig, A., Liu, S., Clasen, L. S., Blumenthal, J. D., Reardon, P. K., Lalonde, F., Greenstein, D., Patel, R., Chakravarty, M. M., Lerch, J. P., & Raznahan, A. (2019). Longitudinally Mapping Childhood Socioeconomic Status Associations with Cortical and Subcortical Morphology. The Journal of Neuroscience, 39(8), 1365–1373. https://doi.org/10.1523/JNEUROSCI.1808-18.2018

McLoyd, V. C. (1998). Socioeconomic disadvantage and child development. The American Psychologist, 53(2), 185–204.

Merz, E. C., He, X., & Noble, K. G. (2018). Anxiety, depression, impulsivity, and brain structure in children and adolescents. NeuroImage : Clinical, 20, 243–251. https://doi.org/10.1016/j.nicl.2018.07.020

Merz, E. C., Maskus, E. A., Melvin, S. A., He, X., & Noble, K. G. (2020). Socioeconomic Disparities in Language Input Are Associated With Children’s Language-Related Brain Structure and Reading Skills. Child Development, 91(3), 846–860. https://doi.org/10.1111/cdev.13239

Merz, E. C., Wiltshire, C. A., & Noble, K. G. (2019). Socioeconomic Inequality and the Developing Brain: Spotlight on Language and Executive Function. Child Development Perspectives, 13(1), 15–20. https://doi.org/10.1111/cdep.12305

Mills, K. L., Goddings, A.-L., Herting, M. M., Meuwese, R., Blakemore, S.-J., Crone, E. A., Dahl, R. E., Güroğlu, B., Raznahan, A., Sowell, E. R., & Tamnes, C. K. (2016). Structural brain development between childhood and adulthood: Convergence across four longitudinal samples. NeuroImage, 141, 273–281. https://doi.org/10.1016/j.neuroimage.2016.07.044

Mills, K. L., & Tamnes, C. K. (2014). Methods and considerations for longitudinal structural brain imaging analysis across development. Developmental Cognitive Neuroscience, 9, 172–190. https://doi.org/10.1016/j.dcn.2014.04.004

Mitchell, B. L., Cuéllar-Partida, G., Grasby, K. L., Campos, A. I., Strike, L. T., Hwang, L.-D., Okbay, A., Thompson, P. M., Medland, S. E., Martin, N. G., Wright, M. J., & Rentería, M. E. (2020). Educational attainment polygenic scores are associated with cortical total surface area and regions important for language and memory. NeuroImage, 212, 116691. https://doi.org/10.1016/j.neuroimage.2020.116691

Noble, K. G., & Giebler, M. A. (2020). The Neuroscience of Socioeconomic Inequality. Current Opinion in Behavioral Sciences, 36, 23–28. https://doi.org/10.1016/j.cobeha.2020.05.007

Noble, K. G., Houston, S. M., Brito, N. H., Bartsch, H., Kan, E., Kuperman, J. M., Akshoomoff, N., Amaral, D. G., Bloss, C. S., Libiger, O., Schork, N. J., Murray, S. S., Casey, B. J., Chang, L., Ernst, T. M., Frazier, J. A., Gruen, J. R., Kennedy, D. N., Zijl, P. V., … Sowell, E. R. (2015). Family Income, Parental Education and Brain Structure in Children and Adolescents. Nature Neuroscience, 18(5), 773–778. https://doi.org/10.1038/nn.3983

Noble, K. G., McCandliss, B. D., & Farah, M. J. (2007). Socioeconomic gradients predict individual differences in neurocognitive abilities. Developmental Science, 10(4), 464–480. https://doi.org/10.1111/j.1467-7687.2007.00600.x

Noble, K. G., Norman, M. F., & Farah, M. J. (2005). Neurocognitive correlates of socioeconomic status in kindergarten children. Developmental Science, 8(1), 74–87. https://doi.org/10.1111/j.1467-7687.2005.00394.x

Okbay, A., Beauchamp, J. P., Fontana, M. A., Lee, J. J., Pers, T. H., Rietveld, C. A., Turley, P., Chen, G.-B., Emilsson, V., Meddens, S. F. W., Oskarsson, S., Pickrell, J. K., Thom, K., Timshel, P., de Vlaming, R., Abdellaoui, A., Ahluwalia, T. S., Bacelis, J., Baumbach, C., … Benjamin, D. J. (2016). Genome-wide association study identifies 74 loci associated with educational attainment. Nature, 533(7604), 539–542. https://doi.org/10.1038/nature17671

Ozernov-Palchik, O., Norton, E. S., Wang, Y., Beach, S. D., Zuk, J., Wolf, M., Gabrieli, J. D. E., & Gaab, N. (2018). The relationship between socioeconomic status and white matter microstructure in pre-reading children: A longitudinal investigation. Human Brain Mapping, 40(3), 741–754. https://doi.org/10.1002/hbm.24407

Pace, A., Luo, R., Hirsh-Pasek, K., & Golinkoff, R. M. (2017). Identifying Pathways Between Socioeconomic Status and Language Development. Annual Review of Linguistics, 3(1), 285–308. https://doi.org/10.1146/annurev-linguistics-011516-034226

Panizzon, M. S., Fennema-Notestine, C., Eyler, L. T., Jernigan, T. L., Prom-Wormley, E., Neale, M., Jacobson, K., Lyons, M. J., Grant, M. D., Franz, C. E., Xian, H., Tsuang, M., Fischl, B., Seidman, L., Dale, A., & Kremen, W. S. (2009). Distinct Genetic Influences on Cortical Surface Area and Cortical Thickness. Cerebral Cortex, 19(11), 2728–2735. https://doi.org/10.1093/cercor/bhp026

Plomin, R., DeFries, J. C., Knopik, V. S., & Neiderhiser, J. M. (2016). Top 10 Replicated Findings From Behavioral Genetics. Perspectives on Psychological Science, 11(1), 3–23. https://doi.org/10.1177/1745691615617439

Plomin, R., & von Stumm, S. (2018). The new genetics of intelligence. Nature Reviews. Genetics, 19(3), 148–159. https://doi.org/10.1038/nrg.2017.104

Price, A. L., Patterson, N. J., Plenge, R. M., Weinblatt, M. E., Shadick, N. A., & Reich, D. (2006). Principal components analysis corrects for stratification in genome-wide association studies. Nature Genetics, 38(8), 904–909. https://doi.org/10.1038/ng1847

Rakic, P. (1988). Specification of cerebral cortical areas. Science, 241(4862), 170–176. https://doi.org/10.1126/science.3291116

Raznahan, A., Shaw, P., Lalonde, F., Stockman, M., Wallace, G. L., Greenstein, D., Clasen, L., Gogtay, N., & Giedd, J. N. (2011). How Does Your Cortex Grow? The Journal of Neuroscience, 31(19), 7174–7177. https://doi.org/10.1523/JNEUROSCI.0054-11.2011

Rea-Sandin, G., Oro, V., Strouse, E., Clifford, S., Wilson, M. N., Shaw, D. S., & Lemery-Chalfant, K. (2021). Educational attainment polygenic score predicts inhibitory control and academic skills in early and middle childhood. Genes, Brain, and Behavior, e12762. https://doi.org/10.1111/gbb.12762

Richardson, J. T. E. (2011). Eta squared and partial eta squared as measures of effect size in educational research. Educational Research Review, 6(2), 135–147. https://doi.org/10.1016/j.edurev.2010.12.001

Rietveld, C. A., Medland, S. E., Derringer, J., Yang, J., Esko, T., Martin, N. W., Westra, H.-J., Shakhbazov, K., Abdellaoui, A., Agrawal, A., Albrecht, E., Alizadeh, B. Z., Amin, N., Barnard, J., Baumeister, S. E., Benke, K. S., Bielak, L. F., Boatman, J. A., Boyle, P. A., … Koellinger, P. D. (2013). GWAS of 126,559 Individuals Identifies Genetic Variants Associated with Educational Attainment. Science, 340(6139), 1467–1471. https://doi.org/10.1126/science.1235488

Rolls, E. T. (2019). The orbitofrontal cortex and emotion in health and disease, including depression. Neuropsychologia, 128, 14–43. https://doi.org/10.1016/j.neuropsychologia.2017.09.021

Romeo, R. R., Christodoulou, J. A., Halverson, K. K., Murtagh, J., Cyr, A. B., Schimmel, C., Chang, P., Hook, P. E., & Gabrieli, J. D. E. (2017). Socioeconomic Status and Reading Disability: Neuroanatomy and Plasticity in Response to Intervention. Cerebral Cortex, 1–16. https://doi.org/10.1093/cercor/bhx131

Selzam, S., Krapohl, E., von Stumm, S., O’Reilly, P. F., Rimfeld, K., Kovas, Y., Dale, P. S., Lee, J. J., & Plomin, R. (2017). Predicting educational achievement from DNA. Molecular Psychiatry, 22(2), 267–272. https://doi.org/10.1038/mp.2016.107

Sherif, T., Rioux, P., Rousseau, M.-E., Kassis, N., Beck, N., Adalat, R., Das, S., Glatard, T., & Evans, A. C. (2014). CBRAIN: A web-based, distributed computing platform for collaborative neuroimaging research. Frontiers in Neuroinformatics, 8, 54. https://doi.org/10.3389/fninf.2014.00054

Skoe, E., Krizman, J., & Kraus, N. (2013). The Impoverished Brain: Disparities in Maternal Education Affect the Neural Response to Sound. The Journal of Neuroscience, 33(44), 17221–17231. https://doi.org/10.1523/JNEUROSCI.2102-13.2013

Smith-Woolley, E., Selzam, S., & Plomin, R. (2019). Polygenic score for educational attainment captures DNA variants shared between personality traits and educational achievement. Journal of Personality and Social Psychology, 117(6), 1145–1163. https://doi.org/10.1037/pspp0000241

Tucker-Drob, E. M., & Bates, T. C. (2016). Large Cross-National Differences in Gene × Socioeconomic Status Interaction on Intelligence. Psychological Science, 27(2), 138–149. https://doi.org/10.1177/0956797615612727

Tulsky, D. S., Carlozzi, N., Chevalier, N., Espy, K., Beaumont, J., & Mungas, D. (2013). NIH Toolbox Cognitive Function Battery (NIHTB-CFB): Measuring Working Memory. Monographs of the Society for Research in Child Development, 78(4), 70–87. https://doi.org/10.1111/mono.12035

van Praag, H., Kempermann, G., & Gage, F. H. (2000). Neural consequences of environmental enrichment. Nature Reviews Neuroscience, 1(3), 191–198. https://doi.org/10.1038/35044558

Vijayakumar, N., Mills, K. L., Alexander-Bloch, A., Tamnes, C. K., & Whittle, S. (2018). Structural brain development: A review of methodological approaches and best practices. Developmental Cognitive Neuroscience, 33, 129–148. https://doi.org/10.1016/j.dcn.2017.11.008

von Stumm, S., Smith-Woolley, E., Ayorech, Z., McMillan, A., Rimfeld, K., Dale, P. S., & Plomin, R. (2020). Predicting educational achievement from genomic measures and socioeconomic status. Developmental Science, 23(3), e12925. https://doi.org/10.1111/desc.12925

Ward, M. E., McMahon, G., Pourcain, B. S., Evans, D. M., Rietveld, C. A., Benjamin, D. J., Koellinger, P. D., Cesarini, D., Consortium, T. S. S. G. A., Smith, G. D., & Timpson, N. J. (2014). Genetic Variation Associated with Differential Educational Attainment in Adults Has Anticipated Associations with School Performance in Children. PLOS ONE, 9(7), e100248. https://doi.org/10.1371/journal.pone.0100248

Wertz, J., Caspi, A., Belsky, D. W., Beckley, A. L., Arseneault, L., Barnes, J. C., Corcoran, D. L., Hogan, S., Houts, R. M., Morgan, N., Odgers, C. L., Prinz, J. A., Sugden, K., Williams, B. S., Poulton, R., & Moffitt, T. E. (2018). Genetics and Crime: Integrating New Genomic Discoveries Into Psychological Research About Antisocial Behavior. Psychological Science, 29(5), 791–803. https://doi.org/10.1177/0956797617744542

Wertz, J., Moffitt, T. E., Agnew-Blais, J., Arseneault, L., Belsky, D. W., Corcoran, D. L., Houts, R., Matthews, T., Prinz, J. A., Richmond-Rakerd, L. S., Sugden, K., Williams, B., & Caspi, A. (2020). Using DNA From Mothers and Children to Study Parental Investment in Children’s Educational Attainment. Child Development, 91(5), 1745–1761. https://doi.org/10.1111/cdev.13329

Winkler, A. M., Kochunov, P., Blangero, J., Almasy, L., Zilles, K., Fox, P. T., Duggirala, R., & Glahn, D. C. (2010). Cortical Thickness or Grey Matter Volume? The Importance of Selecting the Phenotype for Imaging Genetics Studies. NeuroImage, 53(3), 1135–1146. https://doi.org/10.1016/j.neuroimage.2009.12.028

Woo, C.-W., Krishnan, A., & Wager, T. D. (2014). Cluster-extent based thresholding in fMRI analyses: Pitfalls and recommendations. NeuroImage, 91, 412–419. https://doi.org/10.1016/j.neuroimage.2013.12.058

Worsley, K. J., Taylor, J. E., Tomaiuolo, F., & Lerch, J. (2004). Unified univariate and multivariate random field theory. NeuroImage, 23 *Suppl 1*, S189–195. https://doi.org/10.1016/j.neuroimage.2004.07.026

Zelazo, P. D., Anderson, J. E., Richler, J., Wallner-Allen, K., Beaumont, J. L., & Weintraub, S. (2013). II. NIH Toolbox Cognition Battery (CB): Measuring executive function and attention. Monographs of the Society for Research in Child Development, 78(4), 16–33. https://doi.org/10.1111/mono.12032

